# Characterization of West Nile virus Koutango lineage from Phlebotomine Sandflies in Kenya 2021

**DOI:** 10.1101/2024.03.27.587075

**Authors:** Jane Wambui Thiiru, Solomon Langat, Francis Mulwa, Stephanie Cinkovich, Hellen Koka, Santos Yalwala, Samoel Khamadi, Justus Onguso, Nicholas Odemba, Francis Ngere, Jaree Johnson, Timothy Egbo, Eric Garges, Elly Ojwang, Fredrick Eyase

**Affiliations:** Institute for Biotechnology Research, Jomo Kenyatta University of Agriculture and Technology, Nairobi, Kenya; Department of Emerging Infectious Diseases, United States Army Medical Research Directorate-Africa, Nairobi, Kenya; Centre for Virus Research, Kenya Medical Research Institute, Nairobi, Kenya; Global Emerging Infections Surveillance Branch, United States Armed Forces Health Surveillance Division, Maryland, United States; United States Armed Forces Pest Management Board, United States, Silver Spring Maryland D, United States

## Abstract

The West Nile virus (WNV), primarily transmitted by mosquitoes, is one of the most widespread flaviviruses globally, with past outbreaks occurring in the USA and Europe. Recent studies in parts of Africa, including Kenya, have identified the West Nile virus Koutango lineage (WN-KOUTV) among phlebotomine sandfly populations, however, our understanding of this virus remains limited. Hence, this study aimed to characterize WN-KOUTV from phlebotomine sandflies. Sandflies were sampled between 12-16^th^ March 2021 from six villages in Baringo South, Kenya, using CDC light traps. Female sandflies were taxonomically identified and pooled based on genus. Virus isolation was performed in Vero cells. Viral genome was determined using next-generation sequencing. Phylogenetic and molecular clock analyses were done to decipher the virus’s evolutionary relationships. Comparative analyses of amino acid sequences were performed to determine variations. Protein modeling in Pymol was conducted to elucidate variations in key protein regions. Evolutionary pressure analysis investigated the selection pressures on the virus. *In vitro* experiments were done to investigate the virus growth kinetics in mammalian (Vero-E6) and mosquito (C636) cells. We report the isolation of WN-KOUTV from Salabani Baringo South, Kenya. The isolated WN-KOUTV clustered with previously identified WN-KOUTV strains. Comparative analysis revealed unique amino acid at NS5 653. Diversifying pressure was acting NS3 267 of the WN-KOUTV lineage. WN-KOUTV replicates efficiently in Vero-E6 and C636 cells comparable to West Nile virus Lineage 1a, isolated from mosquitoes. The isolation of WN-KOUTV in sandflies points to them as potential vectors, however, vector competence studies would confirm this. The efficient replication in mammalian and mosquito cell lines elucidated its adaptability to host and vector. We speculate the close genetic relationship of WN-KOUTV strains is enabled by the bird migratory route between East and West Africa. If proven, this may point to a potential future pandemic pathway for this virus.

## Introduction

West Nile virus (WNV) is an arthropod-borne virus of the genus *Flavivirus*, family *Flaviviridae*, and a member Japanese Encephalitis serocomplex [1]. It was originally isolated in 1937 from the blood of an adult female in Uganda, during routine surveillance for yellow fever virus [2]. Since its initial emergence, WNV has become the most extensively distributed flavivirus, spanning a vast geographic range that encompasses Africa, the Middle East, Europe, Asia, and the Americas [3]. WNV is maintained in an enzootic transmission cycle involving virus reservoir wild birds, mosquito vectors, as well as final or incidental hosts including humans and horses, which are both dead-end hosts [4]. Although several other mosquito species have been suggested to be vectors of WNV, Culex mosquitoes are the primary competent vectors [5]. Birds of orders Passeriformes and Charadriiformes are highly competent hosts that play a crucial role as carriers, amplifiers, and reservoirs of the virus [6]. Approximately, 80% of human cases infected with WNV typically exhibit no symptoms, while around 20% experience mild flu-like symptoms referred to as West Nile fever (WNF) [7]. A small percentage of the cases, less than 1%, can develop West Nile neuro-invasive disease (WNND), which is a more severe condition characterized by WNV meningitis, encephalitis, or poliomyelitis [8,9]. WNV has caused human and animal infections, and some fatal cases, particularly in America [10] and Europe [11,12]. WNV has been classified into nine lineages (WNV lineage 1 to lineage 9) based on biology, evolution, pathogenicity, and geographic distribution [13]. Except West Nile virus Koutango lineage (WN-KOUTV)-lineage 7, strains of lineages 1 and 2 have exhibited the highest virulence, leading to numerous outbreaks accompanied by severe neurological disease in some cases [11]. In Kenya, West Nile Lineage 1a has been documented through isolation from field-collected mosquitoes [14,15] and *Rhipicephalus pulchellus* ticks [16] as well as through serological evidence [17–19] though epidemics have not been reported. Recently West Nile virus Koutango lineage was isolated from phlebotomine sandflies in Baringo county 2015 GenBank accession numbers: ON158112.1.

WN-KOUTV, predominately found in Africa, was for many years reported as a distinct flavivirus however, based on phylogenetic evidence, it is presently regarded as a distant variant of the WNV [20,21]. WN-KOUTV takes its name from the Koutango district of the Kaolack region in Senegal, where the virus was first recovered from the wild rodent *Tatera kempi* in 1968 [22]. Subsequently, it was detected in *mastomys sp* 1974 in Senegal and later detected in *Lemnyscomys sp.* in Central African Republic [23]. In addition, it has been isolated from a range of arthropods including mosquitoes, ticks, and phlebotomine sandflies [23]. In animal models, investigations done on mice have consistently shown that WN-KOUTV is the most neurovirulent WNV lineage [24–26]. Potential risk for humans has been highlighted following an accidental infection in a Senegalese lab worker characterized by fever and rash [27]. There is also documented serological evidence in humans in Gabon [28] and Sierra Leone [29]. *Culex spp.* primary competent vectors of WNV lineage 1 and 2 were found to be incompetent vectors of WN-KOUTV [30]. However, transovarially transmission has been documented in *Aedes Aegypti* [31]. Also, dissemination of WN-KOUTV by *Aedes aegypti* was demonstrated, but only at high titers [32].

Despite these WN-KOUTV remains a neglected arbovirus with lots of knowledge gaps regarding its persistent, transmission and maintenance cycle, genetic diversity, competent vectors, and hosts. Also, the role of phlebotomine sandflies in its maintenance or transmission is unknown. This study aimed to characterize WN-KOUTV in female phlebotomine sandflies, from study sites in Baringo South, Kenya. The information gained from this study sheds light on the circulation of WN-KOUTV as well as its genetic and *in vitro* characteristics.

## Results

### Sandflies collection and virus isolation

A total of 3,679 female phlebotomine sandflies were collected on March 12-16^th^ 2021 from six villages (Table 1) located within the Baringo South, Kenya (Fig.1). They were organized into 406 pools (≤10 sandflies per pool) according to genus and the locality of collection. Pool BAR/S3/S008, which consisted of *Sergentomyia spp* sandflies trapped in Salabani village (Lat/Lon N0.56187 E36.03038 altitude, 3289 ft) exhibited clear cytopathic effect (CPE). The CPE was characterized by rounding of cells alterations in cell morphology and disruption of the monolayer, starting on day 3 post-inoculation and was consistently observed in the subsequent passages.

**Fig 1:**
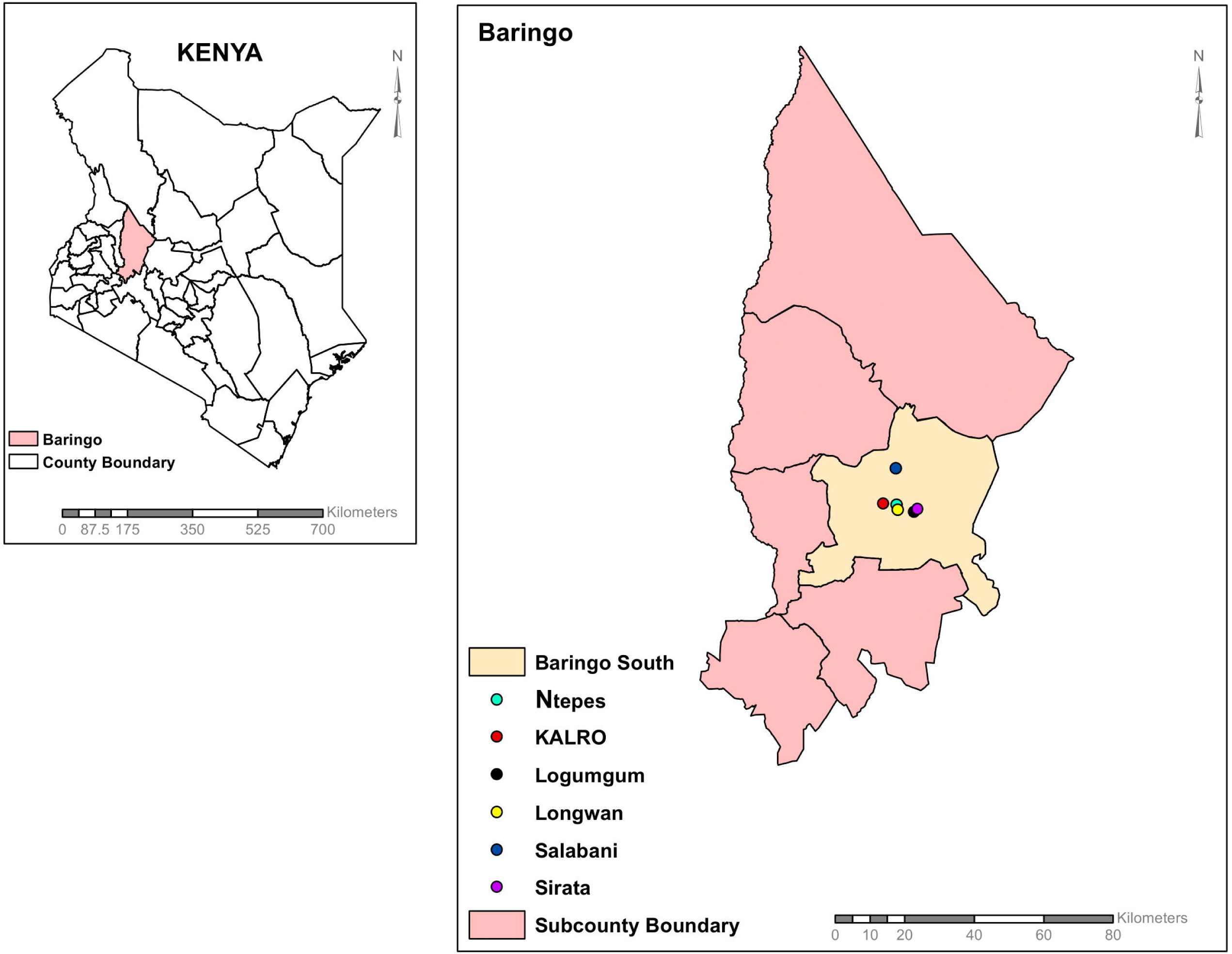
Map of Kenya, Baringo County, and Baringo South showing the sampling sites in this study. The map was developed using ArcGIS Software Version 10.2.2 (http://desktop.arcgis.com/en/arcmap) advanced license.

**Table 1.**
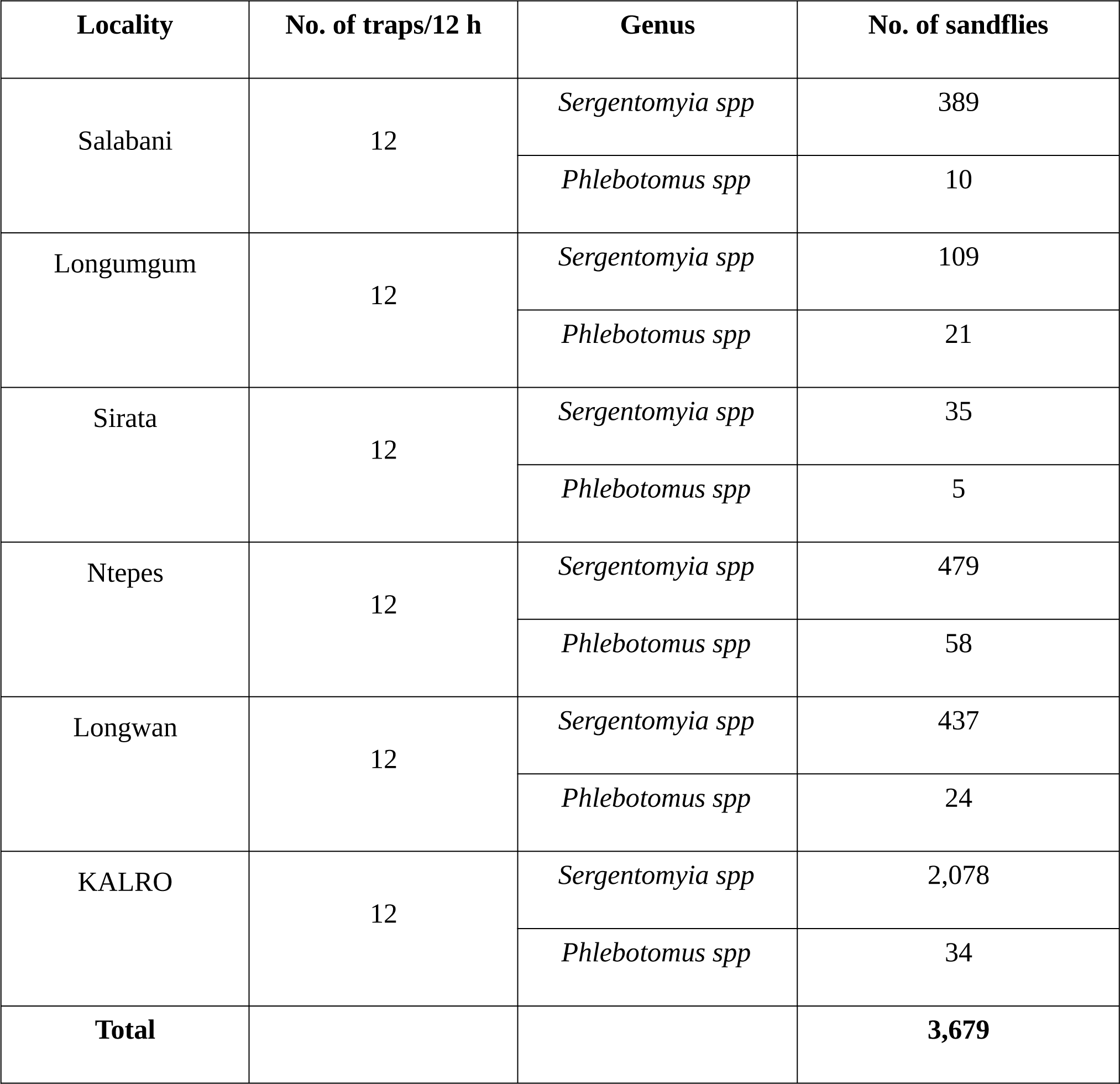
Distribution of phlebotomine sandflies collected from villages in Baringo South using CDC miniature light traps.

### Genetic characterization of the WN-KOUTV isolate

The complete genome of the BAR/S3/S008 isolate was determined. The genome comprises 10,924 nucleotides, with a G+C content of 50.6%. The depth of coverage across the entire genome was well covered, with an average coverage of depth 3864x (S1 Fig). The genome contains a single open reading frame located between 30 and 10,343 nucleotides, encoding a 3437 amino acid long polyprotein. Blast analysis revealed that the genome belongs to a flavivirus of West Nile virus Koutango lineage. Nucleotide similarity to available ORF sequences of other WN-KOUTV strains ranged between 90.11% (PM148; MN057643.1), 90.15% (ArD96655; KY703855.1), 91.20% (DakAnD5443; EU082200.2) and 99.25% (BAR_SS4_020; ON158112.1). At the amino acid level the percent identity ranges from 98.23-98.53 to other WN-KOUTV strains.

### Phylogeny of the WN-KOUTV isolate

Maximum likelihood phylogenetic analysis of the WN-KOUTV isolate, alongside representative strains from various West Nile Virus (WNV) lineages, revealed that the isolate forms a well-supported, single clade with ON158112.1, the WN-KOUTV isolate from Kenya in 2015. This cluster forms a sister clade to the West African WN-KOUTVs, as illustrated in Fig 2.

**Fig 2:**
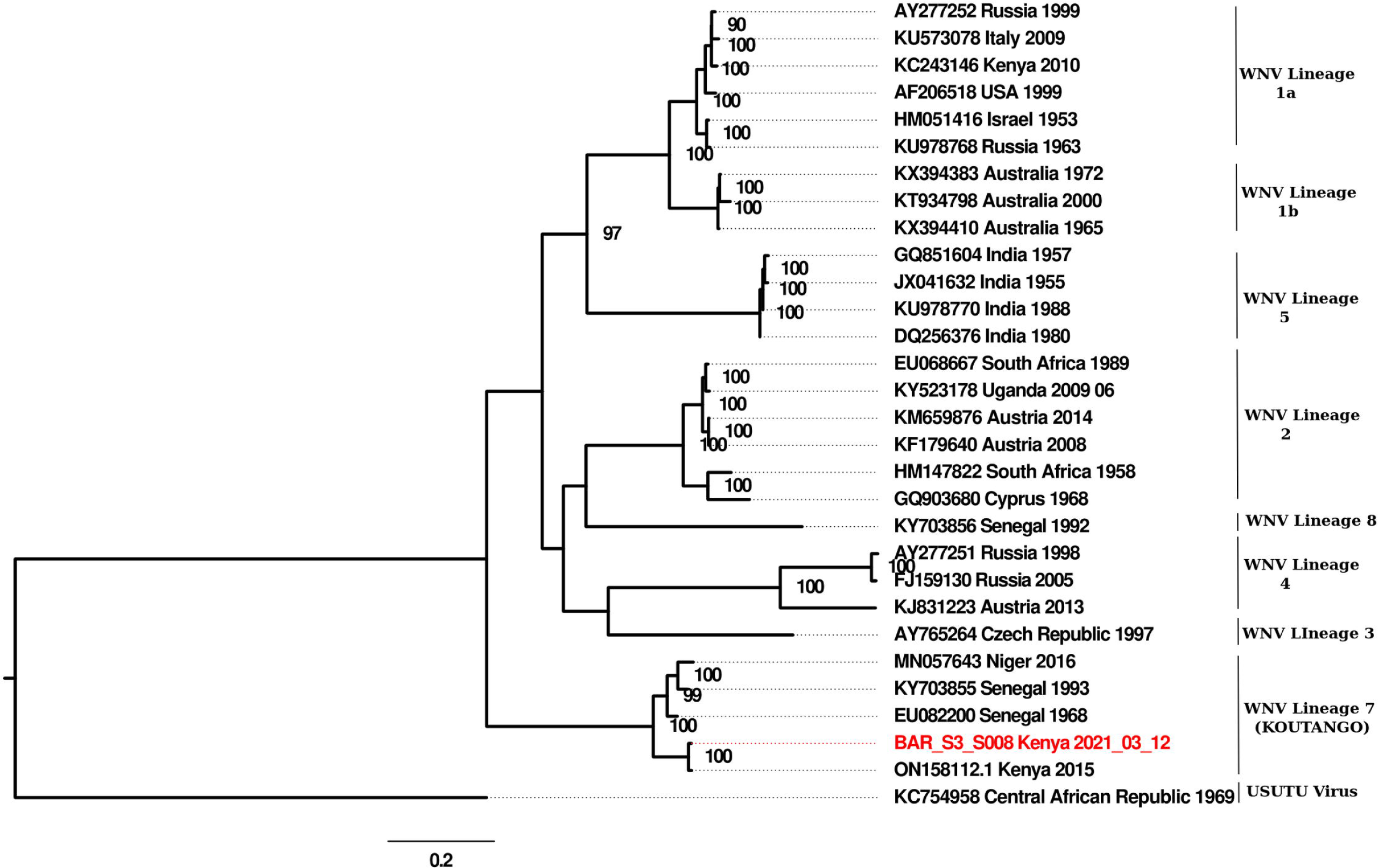
Maximum-likelihood phylogenetic trees showing the relationship of the current WN-KOUTV isolate with select WNV strains. The robustness of the trees was determined using bootstrap analysis. Only bootstrap values >70% are shown at key nodes. Usutu virus was considered as the out-group. The scale bar indicates nucleotide substitutions per site.

The molecular clock analysis estimated the time to the most recent common ancestor (tMRCA) of the WN-KOUTV isolate to be in the early 20th century (1930.91, 95% HPD: 1904.56, 1957.36) (Fig 3) making this the most likely latest time point of introduction of this particular strain in Kenya. The estimated tMRCA of this lineage was dated back to 1885.39.

**Fig 3:**
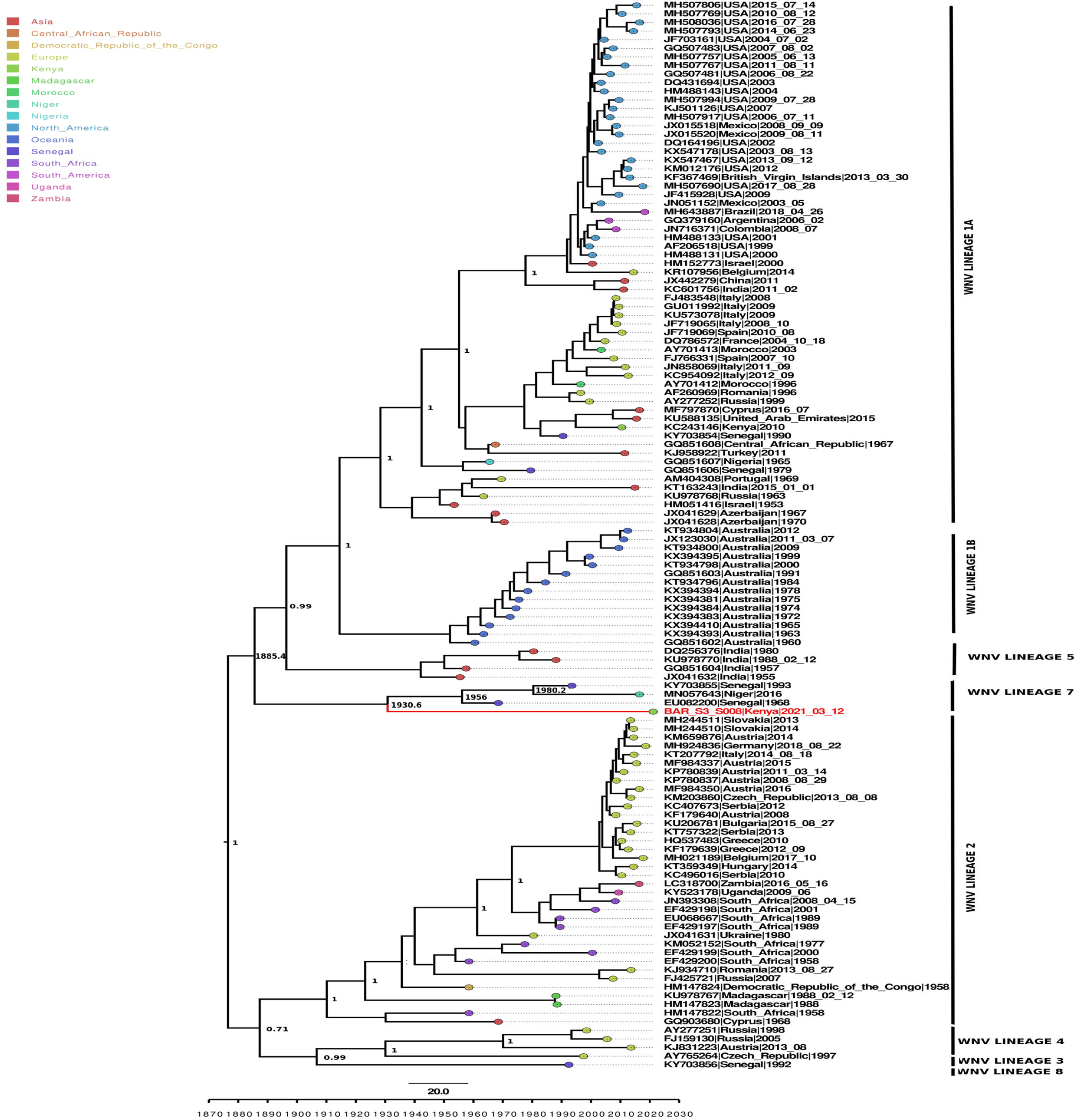
Time-calibrated phylogeny based on 124 genome sequences representing different WNV lineages from different countries. Tree nodes with a posterior probability greater than 0.7 are displayed. The current WN-KOUTV isolate strain is shown in red and tree tip-nodes are colored based on the inferred geographical location of origin for visual clarity. Branches are scaled in years before 2021.28.

### Amino acid variations among WN-KOUTV strains

The amino acid analysis of WN-KOUTV lineage strains’ open reading frame showed 76 out of 3437 (2.21%) variable sites among the strains. The NS2b gene is highly conserved without any variation in the amino acids (Fig 4). The current WN-KOUTV strain showed 41 out of 76 (53.95%) amino acid variations in genes to the other WN-KOUTVs. Amino acid variations were found in the capsid gene 4 (9.75%), envelope gene 6 (14.63), NS1 gene 4 (9.75%), NS2a gene 4 (9.75%), NS3 gene 3 (7.31%), NS4a gene 3(7.31%), NS4b gene 10 (6 amino acid variation and 4 amino acid insertion accounting for 24.39%), and NS5 gene 7 (17.03%) (Fig 4). The insertion of the 4 amino acids at NS4b resulted in a 3437 amino acid-long polyprotein, compared to the 3433-amino acid polyprotein observed in other WN-KOUTV strains. This characteristic is shared with a previous isolate from Kenya, ON158112.1.

**Fig 4:**
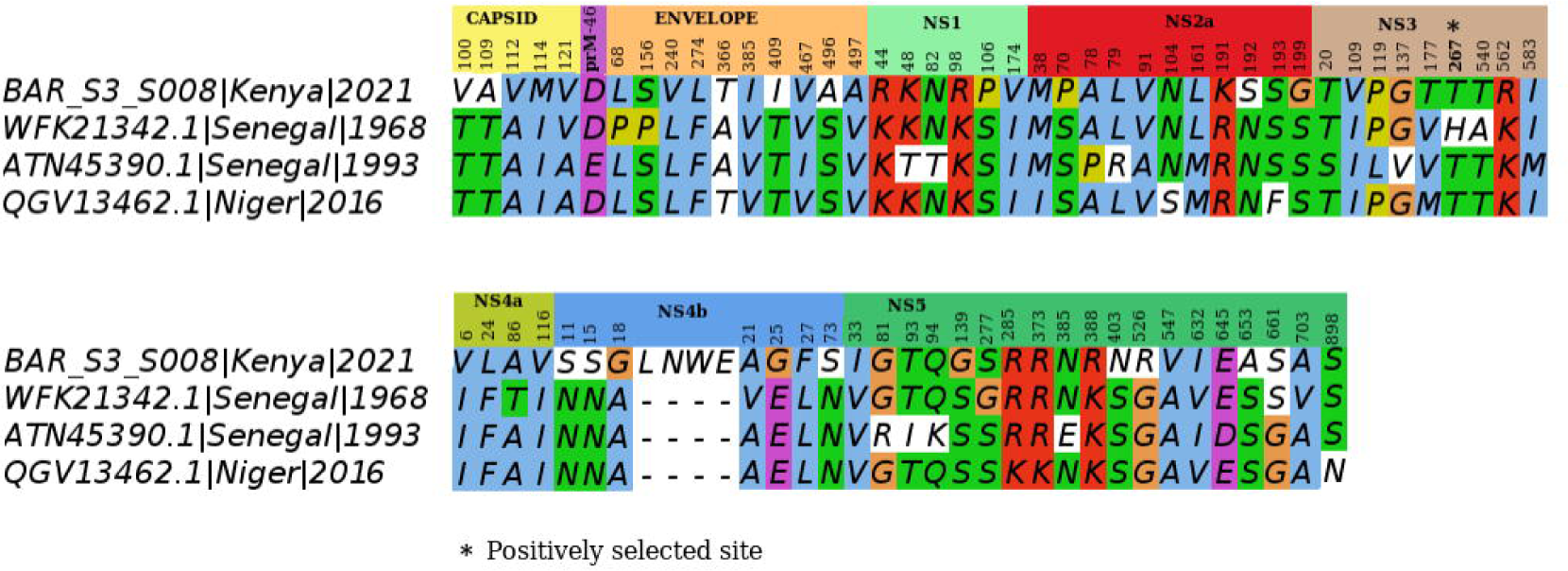
Amino acids variation among the WN-KOUTV strains. Amino acid variations mapped across WN-KOUTV strains polyproteins, revealing gene-specific differences.

Already described mutations characteristic of the WN-KOUTV lineage in published molecular determinants of West Nile Virus virulence and replication were also observed in the current WN-KOUTV isolate, including; (S72M) in the pre-membrane, (Y155F) within the glycosylation site in the envelope and Threonine (T) residue at position 249 of the NS3 protein [6,33,34]. However, a unique amino acid was observed at position 653 in the NS5 protein of the isolated strain. While other WN-KOUTV strains have F653S, the current strain exhibits F653A mutation (Fig 5).

**Fig 5:**
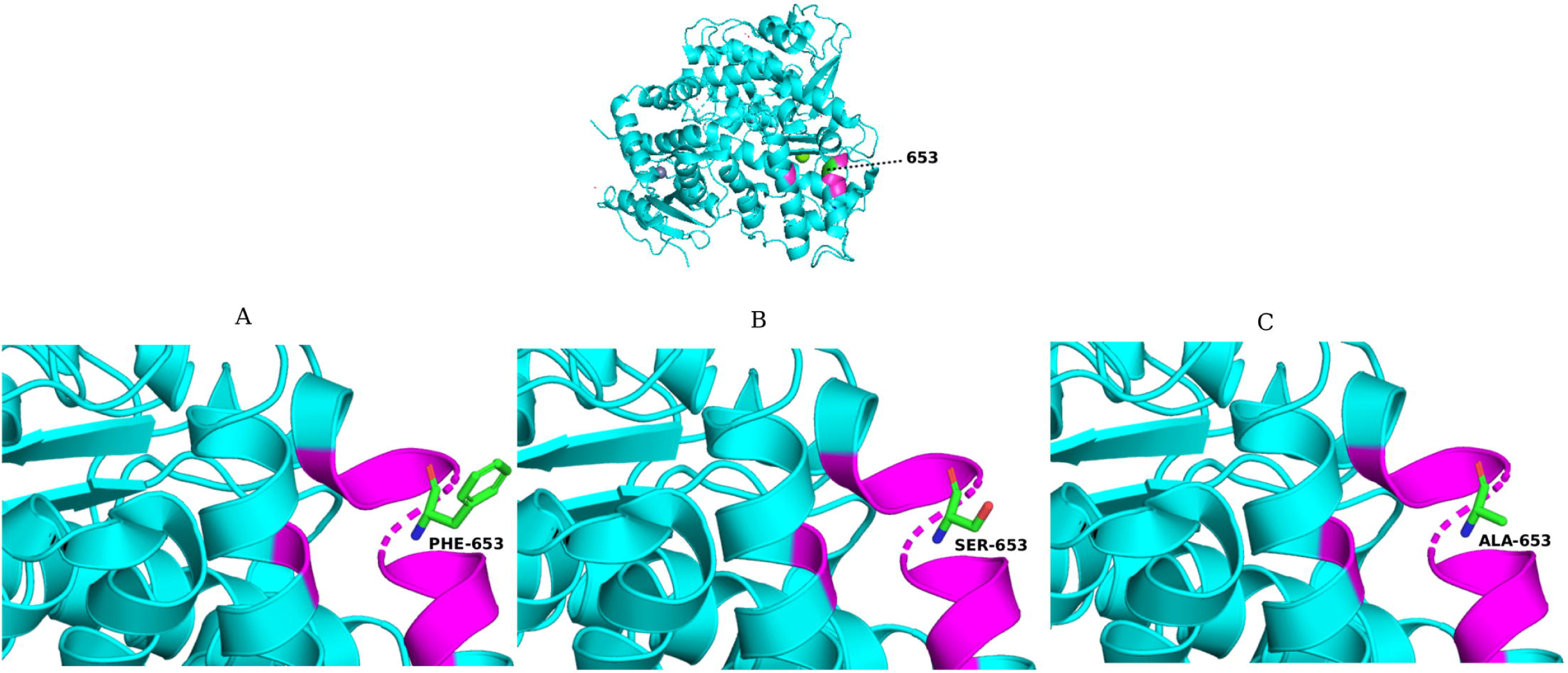
Amino acid variation at position 653 within NS5 Protein. Position of codon 653 (green) and closely associated amino acids (Magenta) mapped on the WNV KUN RdRp structure (Protein Data Bank ID: 2HZF) [35]. A. Phenylalanine Amino acid found in most WNVs lineages strain. Serine mutation found in WN-KOUTVs from West Africa. C. Alanine mutation found in WN-KOUTVs from Kenya.

### Selection pressure was acting at codon site 1772-NS3:267 of WN-KOUTV lineage

A dataset consisting of four WN-KOUTV sequences was assessed for selection pressure. The overall ratio of non-synonymous to synonymous substitutions (dN/dS or ω) was determined to be 0.0216. Significant evidence of positive selection was observed at one site (1772-NS3:267), supported by three of the employed methods namely, FUBAR, FEL, and MEME (Table 2).

**Table 2:**
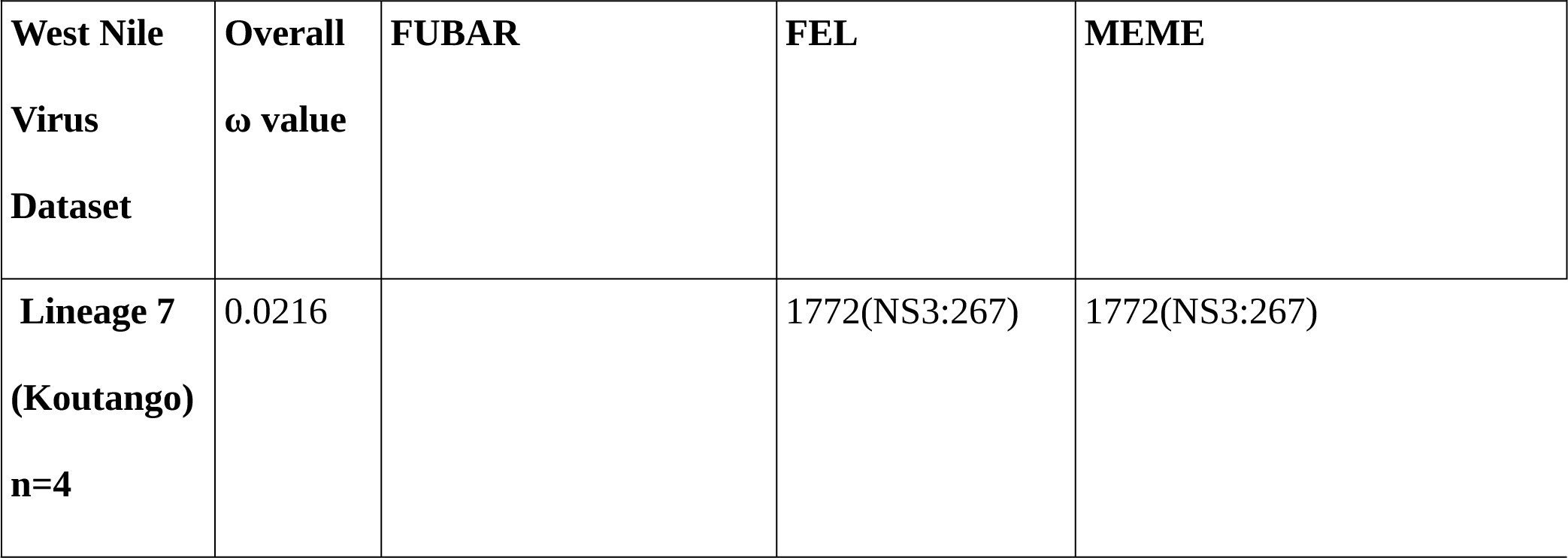
Selection pressure analysis table indicating positively selected site 1772 (NS3:267) of WNV-KOUTV lineage.

### WN-KOUTV replicates efficiently in Vero-E6 and C6/36 cells comparable to WNV Lineage 1a

The WN-KOUTV isolate showed efficient growth in both Vero and C6/36 cells, as indicated by the amount of infectious viral particles (PFU/ml) in the supernatant fraction. In Vero cells, the WN-KOUTV the number of infectious viral particles increased gradually rising to a maximum peak titer 1×10^11^ PFU/mL) 60 hours postinfection (hpi.) before the titers started to decrease. In C6/36 mosquito cells, the WN-KOUTV reached high titers (1×10^10^ PFU/mL) within 144 hours of the experiment. There was no significant difference in infectious viral particle production for the two WNV lineages in Vero cells, however, there was significantly higher rate at 144 hpi for the WN-KOUTV in C6/36 cells (P < 0.0119) (Fig. 6).

**Fig 6:**
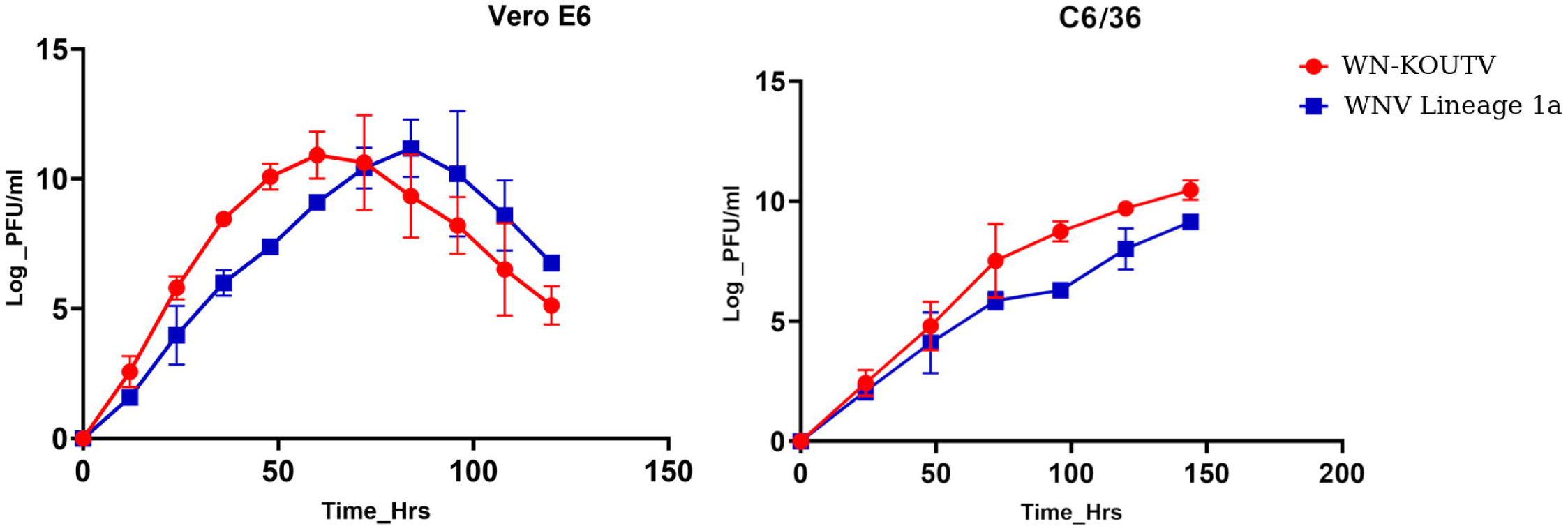
Comparative growth kinetics in WN-KOUTV isolate and WNV Lineage 1a. (A) Vero cells and (B) C6/36 insect cells. Viral titers were determined on Vero E6 cells by plaque assay at the indicated time points. Graphs were plotted in GraphPad Prism 5.02 and statistical significance determined using paired t-test. The error bars indicate the range in values of two independent experiments.

## Discussion

The presence of WN-KOUTV lineage in Africa has been documented through serological evidence, isolation from mammals, and diverse arthropods. Notably, its heightened neurovirulence in animal models compared to other WNV lineages [24,24–26] underscores the need for a deeper understanding of its circulation, genomic traits, and potential vectors. The recent isolation of WN-KOUTV from phlebtomine sandflies highlights their possible role as vectors [23]. The present study successfully isolated and obtained the complete genome of the WN-KOUTV from phlebotomine sandflies sampled from Baringo South, Kenya. This marks the second instance of WN-KOUTV isolation from phlebotomine sandflies in the region, following a previous isolation in 2015 (GenBank accession number: ON158112.1). This subsequent detection of WN-KOUTV in the same region after five years implies ongoing viral circulation and highlights the potential involvement of phlebotomine sandflies, in maintaining the virus within its enzootic transmission cycle. Phlebtomine sandflies in the WN-KOUTV positive pool were morphologically identified as *Sergentomyia spp*. Traditionally, *Sergentomyia spp* are known for their preference to feed on cold-blooded animals [36,37]. However, recent studies have revealed DNA from various vertebrates, including humans, in blood-fed females of different Sergentomyia species, suggesting a potential vector role [38–40]. This isolation also emphasizes the possible wider range of vectors of WN-KOUTV, with previous isolations from mosquitoes and ticks in Senegal [24] and from sandflies in Niger in 2015 [23]

Our molecular clock inference revealed the most recent common ancestor of WN-KOUTV dates to 1930 (95% HPD: 1904.56, 1957.36) (Fig 3) indicative of the possible introduction time of the WN-KOUTV in the region. The extended branch leading to the WN-KOUTV isolate underscores the historical continuity of the virus’s circulation and highlights potential gaps in surveillance data. This may be attributed to the limited attention given to sandfly-borne viruses until recently in the Sub-Saharan region. [40,41]. Historically, surveillance efforts have predominantly focused on vectors such as mosquitoes and ticks [42], potentially allowing the circulation of WN-KOUTV to go undetected for an extended period. Notably, WNV lineage 1a primarily transmitted by mosquitoes, has been documented through serological evidence in this region [19]. Given that birds serve as the natural reservoir hosts for WNVs [6], and considering the documented adaptation of WN-KOUTV to avian cells [43], along with the observed phylogenetic relationship of West and East Africa WN-KOUTV (Fig 3), we speculate that intra-African migratory birds between these regions may have contributed to the movement and potential spread of the virus.

The current WN-KOUTV strain possesses mutations previously observed in West Africa WN-KOUTV strains: S72M in the pre-membrane, Y155F within the glycosylation site in the envelope, and a threonine (T) residue at position 249 of the NS3 protein[23,24,26]. However, it exhibits F653A mutation in the NS5 protein, in place of F653S mutation exhibited by West Africa WN-KOUTV strains (Fig. 5). NS5 653 plays a major role in the resistance to interferon in WNVs [44], hence new studies are necessary to verify possible mutation and its implications.

This study has demonstrated a strong purifying pressure acting on WN-KOUTV (Table 2), aligning with observations in other vector-borne RNA viruses. This strong purifying pressure is attributed to the interaction of the arbovirus with its dual hosts, the arthropod and the vertebrate, each equipped with distinct defense mechanisms [45]. Nevertheless, we found significant diversifying pressure at the codon site 1772 of the polyprotein gene (NS3-267) (Fig 4). NS3-267 is located within the NS3 helicase domain, which is crucial to viral replication [46]. However, no documented role has been attributed to this site in WNV. Therefore, further investigation may elucidate the precise functional implication of this variation.

Our study demonstrates exceptional growth of WN-KOUTV isolate in both mammalian (Vero-E6 cells) and mosquito cell lines (C6/36) comparable to WNV lineage 1a (Fig 6) indicating its adaptability to mammalian and mosquito cell lines despite being isolated from sandflies. Whereas there was no significant difference in infectious particles production between WN-KOUTV and WNV 1a in Vero-E6 cells, there was a significantly higher rate at 144 hpi for the WN-KOUTV in C6/36 cells (p-value 0.0119) in agreement with [24]. Notably, although the WN-KOUTV isolate replicated efficiently in mosquito cells (C6/36), the virus has not yet been isolated from mosquitoes in Kenya. Also, dissemination of DAK

Ar D 5443 WN-KOUTV strain by *Aedes aegypti* was demonstrated at only high virus titers.[32]. Therefore, the re-isolation of WN-KOUTV in phlebotomine sandflies emphasizes the need for further studies to investigate the vector competence of phlebotomine sandflies in the dissemination of WN-KOUTV, as well as serological evidence across avian, human, and other mammalian populations.

## Materials and methods

### Ethics Statement

The study received approval from the Kenya Medical Research Institute (KEMRI) Scientific and Ethics Review Unit under protocols number 3948 and 4570. Additionally, verbal consent was obtained from the heads of households to sample on their farms.

### Study area

The study area encompassed six villages: Longumgum, Salabani, Sirata, Ntepes, Longwan, and KALRO, located in Baringo South sub-county, Baringo County, Kenya (Fig. I). Baringo South is situated in the Rift Valley region of Kenya and is one of the sub-counties within Baringo County. Baringo South covers an area of 1985.11 km^2^ and has a population of approximately 90,388 people according to the 2019 census whose primary economic activity is agro-pastoralism The choice of study areas was informed by the history of febrile illness outbreaks, previous cases, or reports of human fevers of unknown origin, the presence of previously reported arboviruses, including WN-KOUTV, as well as the abundance of phlebotomine sandflies.

### Sandflies sampling

Adult phlebotomine sandflies were sampled using CDC miniature light traps (Model 512, John Hock Co., Gainesville, Florida, USA). Traps were deployed overnight (6 pm-6 am) in favorable habitats, including animal sheds and termite mounds. Captured sandflies were temporarily immobilized using triethylamine, sorted, preserved in liquid nitrogen, and transported to the laboratory at the Kenya Medical Research Institute in Kisumu, where they were stored at –80°C until further processing. All sandflies collected were sorted based on their sex, and specifically, female sandflies were morphologically identified to the genus level using established keys. (Abonnenco, 1951; Kirk & Lewis, 1951). Subsequently, they were pooled in 1.5mL tubes, with a maximum of 10 specimens per pool, based on genus, location, trapping site, and capture day, and stored at −80°C.

### Sandflies homogenization

The sandfly pools were homogenized using a Mini-Beadruptor-16 (Biospec, Bartlesville, OK, USA) in 500 µL of homogenization media (minimum essential media supplemented with 15% fetal Bovine Serum (FBS) (Gibco by Life Technologies, Grand Island, NY, USA), 2% L-glutamine (Sigma, Aldrich), and 2% antibiotic/antimycotic (Gibco by Life Technologies, Grand Island, NY, USA.) with zirconium beads (2.0 mm diameter) for 40 seconds. Subsequently, the homogenate was centrifuged at 2500 rpm for 10 minutes at 4°C using a benchtop centrifuge (Eppendorf, USA), and the supernatant was collected.

### Virus isolation

Virus isolation was performed in Vero (African green monkey kidney) cells (CCL-81™), passage 13, obtained from the Viral Haemorrhagic Fever lab at KEMRI, as described before [47]. Briefly, Vero cells grown overnight at 37 °C and 5% CO_2_ in minimum essential medium supplemented with 2% glutamine, 2% penicillin/streptomycin/amphotericin, 10% fetal bovine serum, and 7.5% NaHCO_3_ in 24-well plates (Corning, Incorporated). At 80% confluence, a 50 µL aliquot of the clarified supernatant from individual pools was inoculated into the wells. The plates were incubated for 1 hour in a humidified incubator at 37 °C and 5% CO_2_ with gentle rocking of the plates side to side every 15 minutes for virus adsorption. Following incubation, 1 mL of maintenance medium, comprising minimum essential medium supplemented with 2% glutamine, 2% penicillin/streptomycin/amphotericin, 2% fetal bovine serum, and 7.5% NaHCO3, was added. The plates were then cultured in a humidified incubator at 37 °C and 5% CO2 and monitored daily for cytopathic effects (CPE) for 14 days [16]. Cultures exhibiting CPE were harvested and further passaged by inoculating them onto fresh monolayers of Vero cells (CCL 81™) in 25-cm^2^ cell culture flasks. After two successive passages, the supernatants of virus-infected vero cell cultures exhibiting cytopathic effect of approximately 70% were harvested from the flasks for virus identification through next-generation sequencing.

### Library preparation and next-generation sequencing

Viral particles from CPE-positive cultures were recovered through 0.22µm filters (Millipore, Merck). From an aliquot of 140 μl of the supernatant, viral RNA was extracted using the QIAamp Viral RNA Mini Kit (Qiagen, Hilden, Germany) and eluted in one step with 60 μl of elution buffer. The extracted nucleic acid was quantified using the Qubit RNA fluorometer (Thermo Fisher Scientific, Waltham, Massachusetts, USA). Sequencing libraries were prepared using Illumina RNA Prep with Enrichment, Tagmentation Kit (Illumina, USA) following the manufacturer’s instructions. The obtained libraries were assessed for quantity using the Qubit 2.0 Fluorometer (ThermoFisher, Waltham, MA, USA). Sequencing was performed on an Illumina Iseq platform (Illumina, San Diego, California, USA) using iSeq 100 i1 Reagent v2 (Illumina, San Diego, California, USA) in a 2 × 150 bp paired-ended configuration.

### Sequence analysis

Raw sequencing data was initially processed for quality using Trimmomatic v0.39 [48]. This step involved the removal of adapters, duplicates, and low-quality sequences. Sequences with phred scores below 30, were used for further analysis. The cleaned reads were then assembled using Megahit v1.2.9 [49] with various k-mer values to reconstruct the viral genome. To identify viral contigs within the assembly, a BLASTn [50] search was conducted against the NCBI databases (https://blast.ncbi.nlm.nih.gov/Blast.cgi). The identified genome was deposited in GenBank under the accession number PP489328. To determine the sequencing depth of the obtained genome, sequencing reads were mapped against the genome using Bowtie2 v2.5.2[51], and the depth of coverage was determined using Samtools depth v1.13.

### Phylogenetic analysis

Complete genomes representing all WNV lineages were retrieved from the Bacterial and Viral Bioinformatics Resource Center (BV-BRC) https://www.bv-brc.org/ and combined with the genome sequence of the isolated virus. Alignment was performed using MAFFT v7.490 [52] and manually curated using BIOEDIT [53]. Phylogenetic analysis was conducted using the Maximum Likelihood method in IQTREE v2.0.7 [54] using GTR+F+I+G4 (General Time Reversible + Frequencies + Invariant Sites + Gamma distribution with 4 rate categories) substitution model as determined using Model-finder [55]. The robustness of the tree topology was assessed using 1000 non-parametric bootstrap analyses [56]. The resulting tree was annotated and visualized in FigTree v.1.4.4 (available from http://tree.bio.ed.ac.uk/software/figtree/)

### Molecular clock analysis

To estimate coalescent times of most recent common ancestor (tMRCA) for the WN-KOUTV isolate, molecular clock analysis was performed using the Bayesian Markov Chain Monte Carlo (MCMC) method implemented in BEAST v2.1.3 software package [57]. Complete genomes with country and year of isolation data, from all WNV lineages spanning continents worldwide were obtained from the Bacterial and Viral Bioinformatics Resource Center (BV-BRC) https://www.bv-brc.org/. These genomes were added to the dataset for maximum likelihood phylogenetic analysis, resulting in a total of (n=124) genomes with an alignment length of 11,302 base pairs. The analysis employed the uncorrelated lognormal relaxed molecular clock model to account for rate variation across branches in the phylogenetic tree. Geographical traits were assigned to the sequences, and a flexible non-parametric Bayesian Skyline model and an asymmetric trait model were used to estimate past population dynamics and ancestral states, respectively. The MCMC analyses were run for 800 million states and sampled every 100,000th generation. Tracer v1.7.1 [58] was employed to assess the logs and ensure adequate effective sample sizes (ESS) values > 200 by visual inspection of chains. Maximum clade credibility (MCC) trees were generated using TreeAnnotator v2.1.2, discarding the initial 10% as burnin. The resultant phylogeny was visualized using FigTree v.1.4.4 (available from http://tree.bio.ed.ac.uk/software/figtree/) showing posterior probability values in each node and the time to the most recent common ancestors (tMRCA) as median year with 95% Highest Posterior Density (HPD).

### Amino acid variations analyses and computational protein modeling

Comparative amino acid analyses were performed using a dataset containing 4 WN-KOUTV complete coding regions. Amino acid alignment was performed using MEGA11 [59], showing only variable sites in each WN-KOUTV gene. The unique amino acid variation at the site of interest was mapped onto the Crystal structure of RNA-dependent RNA polymerase domain from West Nile virus (Protein Data Bank ID: 2HZF) [35] using the PyMOL Molecular Graphics System, Version 2.6 Schrödinger, LLC.

### WN-KOUTV lineage selection pressure analyses

Evolutionary pressures acting across the entire coding sequence of WN-KOUTV lineage were assessed using various methods, including fixed-effects likelihood (FEL), single-likelihood ancestor counting (SLAC), mixed effects model of evolution (MEME), and fast unconstrained Bayesian approximation (FUBAR) on the Datamonkey selective and evolutionary bioinformatics online server [60]. To determine the selective pressure, the ratio (ω) of the rate of non-synonymous substitutions (dN) to the rate of synonymous substitutions (dS) per codon site was calculated using the SLAC and FEL codon-based maximum likelihood approaches. A value of ω ≥ 1 indicates positive selection, ω ≤ 1 indicates negative selection, and ω = 0 indicates neutral selection. For statistical significance, the following thresholds were applied: p < 0.05 for SLAC, FEL, and MEME, and a posterior probability of ≥ 0.9 for FUBAR. Positive selection at a specific site was considered present when at least three methods detected it.

### Time course infection of Vero E6 and C6/36 cells with WN-KOUTV isolate in comparison to WNV Lineage 1a

To compare WN-KOUTV isolate with West Nile Lineage 1a growth kinetics in mammalian and mosquito cell lines. Time course infection of Vero E6 (Ceropithecus aethiops) cells and C6/36 cells (*<u>Aedes albopictus</u>*) was carried out as previously described [24]. Briefly, virus stocks of the WN-KOUTV isolate and the previously isolated Kenyan WN Lineage 1a strain (AMH005348) were generated from passage 3 in Vero E6 (Ceropithecus aethiops) cells (Sigma Aldrich, France) and used in this assay. The viral stocks titers were determined using the Vero plaque assay, resulting in titers of 1.1×10^6^ and 1×10^6^ PFU (plaque-forming units) for WN-KOUTV and WNV lineage 1a, respectively. 2×10^5^ Vero E6 cells and C6/36 cells were seeded in 24-well plates. At 80% confluency, cells were infected with 2×10^3^ PFU of the virus in 50 μl of maintenance medium, for a multiplicity of infection (MOI) of 0.01. After a 1-hour incubation period for virus adsorption, minimum essential medium supplemented with 2% glutamine, 2% penicillin/streptomycin/amphotericin, 2% fetal bovine serum, and 7.5% NaHCO_3_ was added, and this time point was set as the starting point for the growth curves (T0). The cells were then incubated in a humidified incubator at 37 °C and 5% CO2 for Vero cells and at 28 °C for C6/36 cells. The infectious supernatant was harvested every 12 hours for 5 days for Vero cells and every 24 hours for C636 cells for 6 days. All collected samples were stored at −80 °C, and subsequent viral titers were determined using the plaque assay method. Two independent experiments were performed to determine the reproducibility of the results.

### Infectious virus quantification

Infectious WN-KOUTV and WNV lineage 1a were quantified via plaque assay. Briefly, 2.3×10^6^ Vero E6 cells were seeded in 12 well plates a day prior to the infection. Supernatants were serially diluted 1:10 in maintenance media before inoculation of 100ul into 75-95% confluent cells. Plates were incubated at 37 °C and 5% CO2 for 1 h for virus adsorption with rocking back and forth every 15 mins. Following the adsorption process, 2.5% methylcellulose overlay was added to each well, and plates were incubated at 37 °C and 5% CO2 for six days. Methylcellulose was aspirated out and cells were fixed with 3.7%v/v formaldehyde (Sigma) overnight, followed by staining with 0.5% crystal violet (Sigma) diluted in absolute ethanol. The plate were then soaked in tap water for 10 minutes to remove excess stain and allowed to dry followed by manual counting of plaques.

### Statistical analysis

Paired t-test analyses were conducted in Graph Pad Prism v*10.1.2* (GraphPad, San Diego, CA, USA, 2016) to determine significant difference in infectious viral particles production between WN-KOUTV and WNV lineage 1a within the same cell line, with a significance threshold of p < 0.05.

### Conclusion

In conclusion, our study highlights the presence, genomic characteristics, and *in vitro* growth kinetics of WN-KOUTV in Baringo South, Kenya. The isolation from sandflies suggests their potential role as vectors, while its adaptability to mammalian and mosquito cell lines provides valuable insights into potential hosts and transmission dynamics. This finding emphasizes the need for enhanced surveillance efforts in other regions of Africa to gauge the presence and potential spread of WN-KOUTV. Further characterization of the virus will contribute to a better understanding of its behavior and inform strategies for prevention and control.

## Acknowledgment

We thank David Oullo, Charles Waga, Richard Ochieng, Eunice Achieng, Daniel Ngonga and Vitalice Opondo for their expert contribution in sandfly sampling, processing and identification. Special thanks to Samuel Owaka for his skillful generation of the map. We are also grateful to the Viral Hemorrhagic Fever Laboratory at KEMRI for generously providing the West Nile Lineage 1 virus isolate.

