## Supplementary figures and images for "Characterization of West Nile virus Koutango lineage from Phlebotomine Sandflies in Kenya 2021"

### S1 Fig

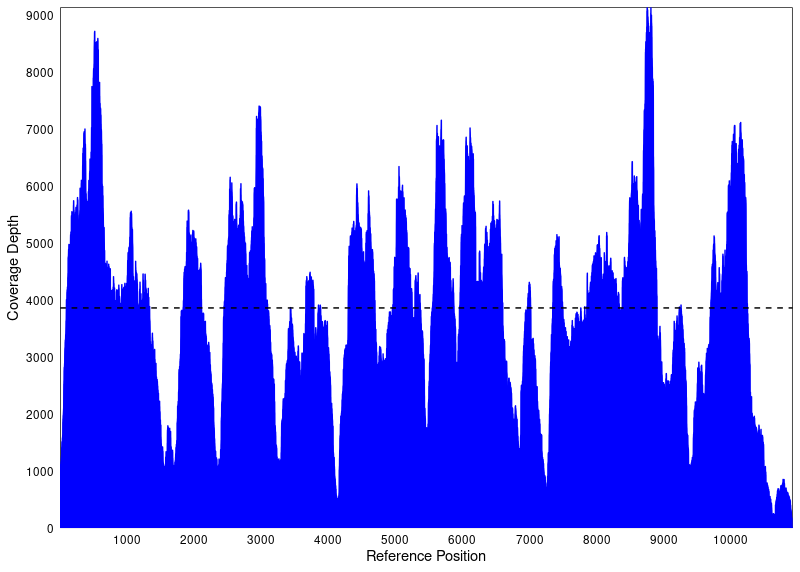
